# Spatially Integrated Multi-Omics reveals the Multicellular Landscape of Progenitor-Driven Glioblastoma Progression

**DOI:** 10.64898/2026.02.26.708154

**Authors:** Korbinian Traeuble, Jakob Träuble, MOSAIC consortium, Gabriele S. Kaminski Schierle, Matthias Heinig

## Abstract

Glioblastoma is the most lethal primary brain tumor, driven by complex interactions between plastic malignant cells and a diverse tumor microenvironment. Despite advances in single-cell profiling, how genomic drivers and the tumor microenvironment interact to determine tumor progression and patient survival remains poorly understood. While cellular states have been cataloged, the multi-cellular logic coordinating these into lethal phenotypes remains unresolved. Here, we integrate whole-exome sequencing, bulk and single-cell RNA sequencing, spatial transcriptomics, and histopathology from the MOSAIC cohort (*n* = 89) to deconstruct this inter- and intra-tumoral heterogeneity. We identify a robust latent multi-omic program in glioblastoma that delineates a transition from homeostatic neural precursors to an aggressive, immunosuppressive progenitor phenotype. This high-risk state, which predicts poor survival in both the MOSAIC discovery and independent validation cohorts (TCGA, CGGA; total *n* = 598), is sustained by dense intercellular communication networks linking malignant progenitors with myeloid and endothelial compartments. Spatially, this program maps to hypoxic, perinecrotic niches, directly linking molecular signaling with microvascular proliferation and tissue necrosis. Our findings provide a spatially resolved, multi-omic blueprint of the multicellular logic driving glioblastoma progression, offering a robust molecular framework for patient stratification and targeted intervention.

## Introduction

Glioblastoma (GBM) represents the most aggressive and lethal primary brain tumor in adults ^1^. Despite a maximalist therapeutic regimen comprising surgical resection followed by concurrent radiochemotherapy with temozolomide, the disease remains essentially incurable, with a median survival plateauing at approximately 12–15 months ^2^. Beyond the standard clinical protocol, *MGMT* promoter methylation is a primary prognostic biomarker; however, its clinical utility is frequently challenged by discordance between methylation status and protein expression levels, suggesting that more comprehensive molecular profiling is required to accurately predict response to alkylating agents ^3,4^. A central obstacle in the clinical management of GBM is its extensive cellular and molecular heterogeneity, which manifests both inter-tumorally and intra-tumorally. Early transcriptomic studies utilizing bulk sequencing established a framework of discrete transcriptional subtypes, Proneural, Classical, and Mesenchymal, driven by distinct genomic alterations such as *PDGFRA* amplification, *EGFR* amplification, and *NF1* loss, respectively ^5^. However, the rigidity of this classification has been challenged by the recognition that GBM tumors are dynamic ecosystems where malignant cells exhibit significant plasticity ^6^.

Recent single-cell RNA sequencing (scRNA-seq) analyses have refined this view, demonstrating that rather than adhering to fixed lineages, GBM cells traverse a continuous landscape of transcriptional states. These states range from neural progenitor-like (NPC) and oligodendrocyte progenitor-like (OPC) identities to astrocyte-like (AC) and mesenchymal-like (MES) states, often influenced by the tumor microenvironment^7^. This plasticity is hierarchically organized, with developmental gradients often mirroring injury-response programs that fuel therapeutic resistance ^8^. Furthermore, it has been suggested that GBM recapitulates a normal neurodevelopmental hierarchy, with a tri-lineage architecture centered on undifferentiated progenitor cells that fuel tumor growth ^9^. This complex neoplastic biology is sustained by a diverse and immunosuppressive tumor microenvironment (TME) ^10^. The GBM TME contains a mix of differentiated cancer cells, therapy-resistant glioma stem-like cells (GSCs) that drive recurrence ^11^, and a variety of stromal and immune cells. In particular, immunosuppressive components, including tumor-associated macrophages (TAMs), myeloid-derived suppressor cells (MDSCs), and regulatory T cells, create a “cold” immune niche that sustains GBM growth and blunts the effectiveness of emerging immunotherapies ^12,13^.

To navigate this cellular complexity, the field has increasingly relied on large-scale multi-modal reference atlases. Foundational resources such as the Ivy Glioblastoma Atlas Project (Ivy GAP) ^14^ have mapped transcriptomic features to specific anatomic niches (e.g., cellular tumor, leading edge, microvascular proliferation), providing a spatial scaffold for understanding tumor architecture. More recently, the integration of diverse single-cell datasets into harmonized references, such as GBmap ^15^, has enabled the robust identification of rare cell types and the deconvolution of bulk profiles, establishing a comprehensive taxonomy of GBM cellular diversity. Complementing these efforts, the recently compiled GRIT-Atlas ^16^, comprising nearly one million cells, has unveiled a treatment-induced fibrotic niche that physically shields recurrent tumors from immunotherapy. The utility of such integrative high-resolution atlases extends beyond oncology; similar efforts have recently expanded our understanding of other inflammation-driven pathologies ^17^. In the context of GBM, these atlases provide the molecular definitions required to resolve tissue heterogeneity, allowing researchers to pinpoint how specific spatial neighborhoods drive malignancy.

Building on this cellular framework, recent advances in spatial transcriptomics have begun to disentangle tumor architecture with unprecedented resolution ^18^. For example, spatial profiling has shown that stem-like glioma cells and specific myeloid populations colocalize in hypoxic pseudopalisading regions, where reciprocal signaling promotes tumor growth and immune suppression ^19^. Similarly, glioblastoma-derived cytokines can reprogram astrocytes into an immunosuppressive state characterized by TRAIL expression, which induces apoptosis in infiltrating T cells and accelerates recurrence ^20^. Such studies highlight that interactions between malignant and non-malignant cells are central to GBM progression.

To capture these interactions across multiple molecular layers, integrative multi-omics approaches are required ^21^. The biology of GBM spans genomic alterations, gene expression programs, epigenetic states, and tissue architecture, all of which contribute to disease course. Multi-Omics Factor Analysis (MOFA) ^22^ provides an unsupervised and interpretable framework for jointly analyzing heterogeneous data types and identifying latent factors that represent the major biological axes of variation. MOFA has been successfully applied in other disease contexts, including chronic lymphocytic leukemia ^23^ and cardiovascular syndromes ^24^, to uncover multicellular programs predictive of clinical status.

Here, we apply MOFA to the Multi-Omics Spatial Atlas in Cancer (MOSAIC) cohort ^25^, a large-scale multimodal resource of 89 GBM samples profiled by whole-exome sequencing, bulk RNA sequencing, scRNA-seq, spatial transcriptomics, and histopathology. By integrating these diverse modalities, we identify a coordinated multicellular program that delineates a transition from homeostatic neural precursors to an aggressive, immune-suppressive progenitor phenotype. We demonstrate that this this progenitor-driven high-risk state is physically mapped to hypoxic, perinecrotic niches and is driven by specific ligand–receptor circuits that connect malignant progenitors with myeloid and endothelial populations. Our results provide a spatially resolved, multi-omic blueprint of the cellular circuits driving progression and offer a robust molecular framework for patient stratification.

## Results

### A multi-modal atlas of glioblastoma heterogeneity

To systematically dissect the molecular and cellular heterogeneity of glioblastoma, we leveraged the multi-modal data of the MOSAIC cohort, which comprises *n* = 89 patients with newly diagnosed, IDH-wildtype glioblastoma, for whom matched whole-exome sequencing (WES), bulk RNA sequencing, single-cell RNA sequencing (scRNA-seq), spatial transcriptomics, and histopathological profiling were available (Fig. 1A). The median patient age at diagnosis was 65.8 years (range: 37.1–89.2), with a sex distribution of 64.0% male and 36.0% female. Molecular stratification indicated that 57.7% of tumors were MGMT promoter unmethylated, while 42.3% were methylated.

**Figure 1:**
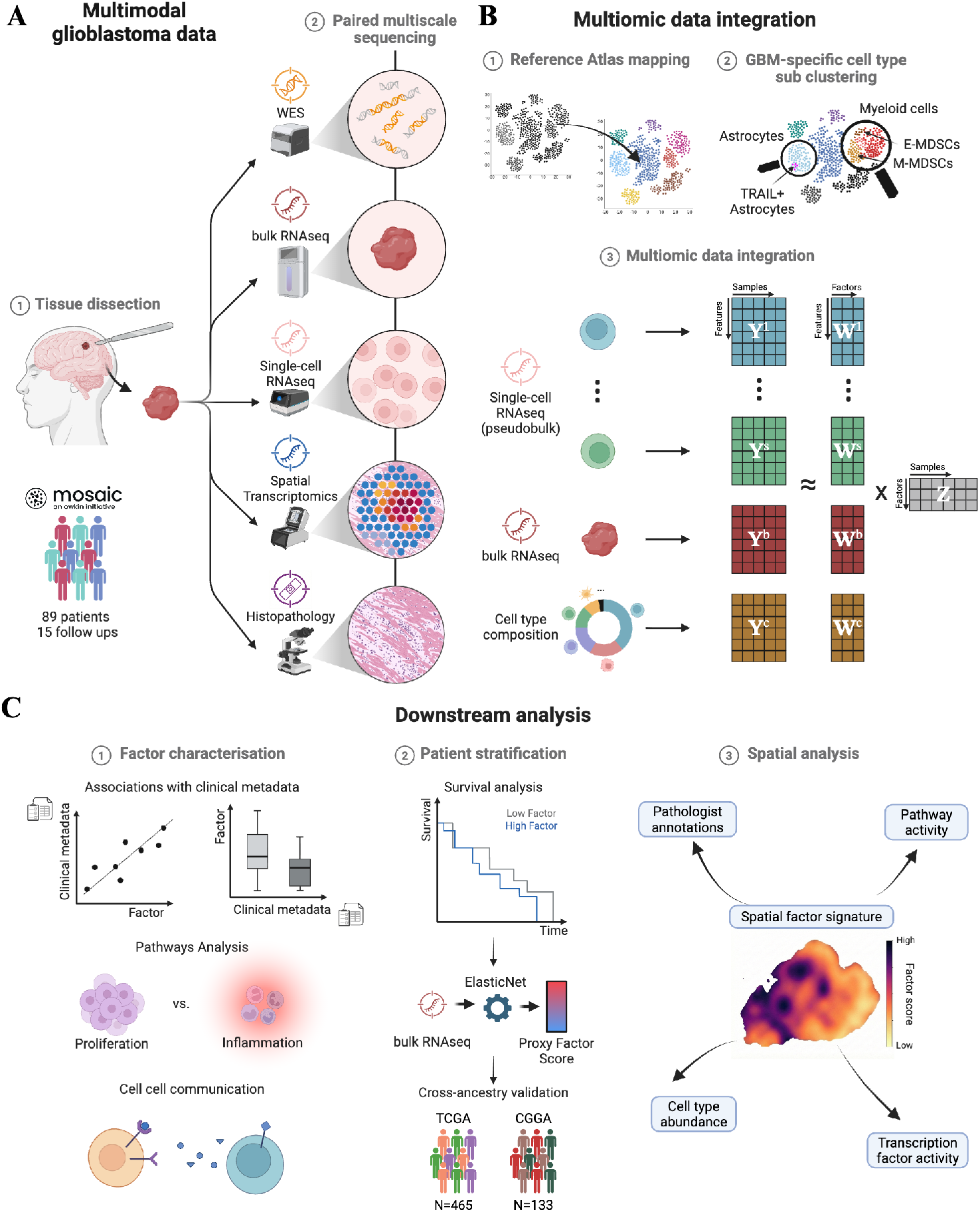
A multi-scale multi-omics approach to glioblastoma heterogeneity. **A**, Study design and data acquisition. Tumor specimens from the MOSAIC cohort (*n* = 89 patients, with 15 paired longitudinal samples) were profiled using whole-exome sequencing (WES), bulk RNA-seq, single-cell RNA-seq (scRNA-seq), spatial transcriptomics, and histopathology. **B**, Data integration strategy. scRNA-seq data were mapped to a reference atlas and sub-clustered to resolve granular GBM-specific cell states. These features (pseudobulk expression, cell type composition) were integrated with bulk RNA-seq using Multi-Omics Factor Analysis (MOFA) to decompose variance into latent factors. **C**, Downstream analysis workflow. Identified factors were characterized via associations with clinical metadata, biological pathways, and cell–cell communication networks. Patients were stratified based on factor values for suvival analysis, and a proxy bulk RNA-seq signature was developed using regression modeling to validate findings in independent cohorts (The Cancer Genome Atlas (TCGA), *n* = 465; Chinese Glioma Genome Atlas (CGGA), *n* = 133). Finally, spatial transcriptomics data were leveraged to generate spatial factor signatures, which were spatially correlated with histopathological annotations, pathway and transcription factor activities, and estimated cell type abundances. Created in https://BioRender.com.

To resolve the cellular heterogeneity of these tumors at high resolution, we utilized the integrative computational framework MOFA (Fig. 1B). This unsupervised approach identifies major axes of variation across multiple data ‘views’, enabling the discovery of latent factors that uncover relationships between molecular layers. To integrate the single-cell transcriptomics, the data was pseudobulked per cell type to account for varying cell numbers while retaining cell-type-specific information. We employed transfer learning to map the cohort’s scRNA-seq data onto a comprehensive reference atlas of over one million glioma cells ^15^, enabling the robust identification of 21 distinct cell types including canonical malignant states (AC-like, NPC-like, OPC-like, MES-like) and diverse immune and stromal populations. Targeted sub-clustering further resolved granular immunosuppressive subsets, such as monocytic and granulocytic myeloid-derived suppressor cells (M-MDSCs, E-MDSCs) and TRAIL+ astrocytes. We integrated this high-resolution cellular landscape with bulk transcriptomics using MOFA to decompose inter-patient variability into latent factors representing co-varying cellular and molecular programs.

To translate these latent dimensions into mechanistic and clinical insights, we devised a multi-stage downstream analysis workflow (Fig. 1C). Identified factors were first functionally annotated through associations with clinical metadata, biological pathway activities, and inferred cell–cell communication networks. We then evaluated their prognostic value using survival analysis and developed a transcriptomic proxy signature, trained via elastic net regression, to validate these risk profiles in independent large-scale cohorts (The Cancer Genome Atlas (TCGA)^26,27^, *n* = 465; Chinese Glioma Genome Atlas (CGGA) ^28,29,30,31^, *n* = 133). Finally, we spatially resolved these programs by projecting factor weights onto matched spatial transcriptomics sections, allowing us to directly correlate latent molecular factors with histopathological features, local pathway activity, and transcription factor regulation in situ.

### The genomic and single-cell transcriptomic landscape of the glioblastoma cohort

We first characterized the genomic alteration landscape of the cohort using WES, specifically focusing on established GBM driver genes. Single-nucleotide variants (SNVs) and copy-number variants (CNVs) were restricted to genes identified as significantly altered in the TCGA Glioblastoma study ^27^ (see Methods). Across the cohort, frequently occurring alterations predominantly affected canonical GBM drivers and core oncogenic pathways (Fig. 2A). The most frequently mutated gene was *TP53*, harboring oncogenic mutations in 33.7% of tumors, followed by *PTEN* (31.5%) and *EGFR* (23.6%). Additional common SNVs were observed in *PIK3CA* (14.6%), *NF1* (13.5%), and *PIK3R1* (6.7%), highlighting frequent disruption of the PI3K/AKT and RAS/MAPK signaling pathways. Together, the SNV landscape underscores widespread inactivation of tumor suppressors alongside alterations in receptor tyrosine kinase and downstream signaling pathways. These prevalence rates align closely with those reported in the TCGA discovery cohort ^27^, confirming that our dataset is representative of the wider IDH-wildtype glioblastoma population. Copy-number analysis revealed a complementary pattern of genomic dysregulation dominated by focal amplifications and deletions affecting established GBM driver loci. The most frequent amplification involved *CDK6* (52.8%), followed by *FSTL3* (33.7%), *ARID3A* (29.2%), *SOX2* (28.1%), and *PDGFRA* (18.0%). In parallel, frequent deletions were most prominent at the CDKN2A/CDKN2B locus, with homozygous loss observed in 76.4% of tumors, accompanied by frequent deletions of *MTAP* (60.7%), *ADARB2* (32.6%), *PTEN* (15.7%), and *RB1* (12.4%). These CNV patterns emphasize hallmark GBM mechanisms of oncogene activation through amplification of cell-cycle and growth-factor signaling genes, together with inactivation of tumor suppressor pathways governing cell-cycle control and genomic stability (Fig. 2A). Combined genetic alterations of the restricted gene set shows *EGFR* as the most alterated gene with 67% alterations (see Supplementary Fig. 3). To investigate tumor evolution, paired primary–relapse WES data were available for 16 patients. Focusing on the most frequently altered genes, *TP53* and *EGFR* are the most frequently changing SNVs across pairs, while *FSTL3* and *ARID3A* showed the strongest relapse-associated CNV amplifications, and *HMGA2* and *RB1* were the most frequently altered CNV deletions (Fig. 2B). Notably, gains and losses were observed across different patients for the same genes, indicating heterogeneous evolutionary trajectories rather than a single dominant relapse program. This observation is consistent with large-scale longitudinal analyses by the GLASS consortium, which demonstrated that glioblastoma recurrence is typically driven by branched evolution and patient-specific genetic divergence rather than a uniform linear progression ^32^.

**Figure 2:**
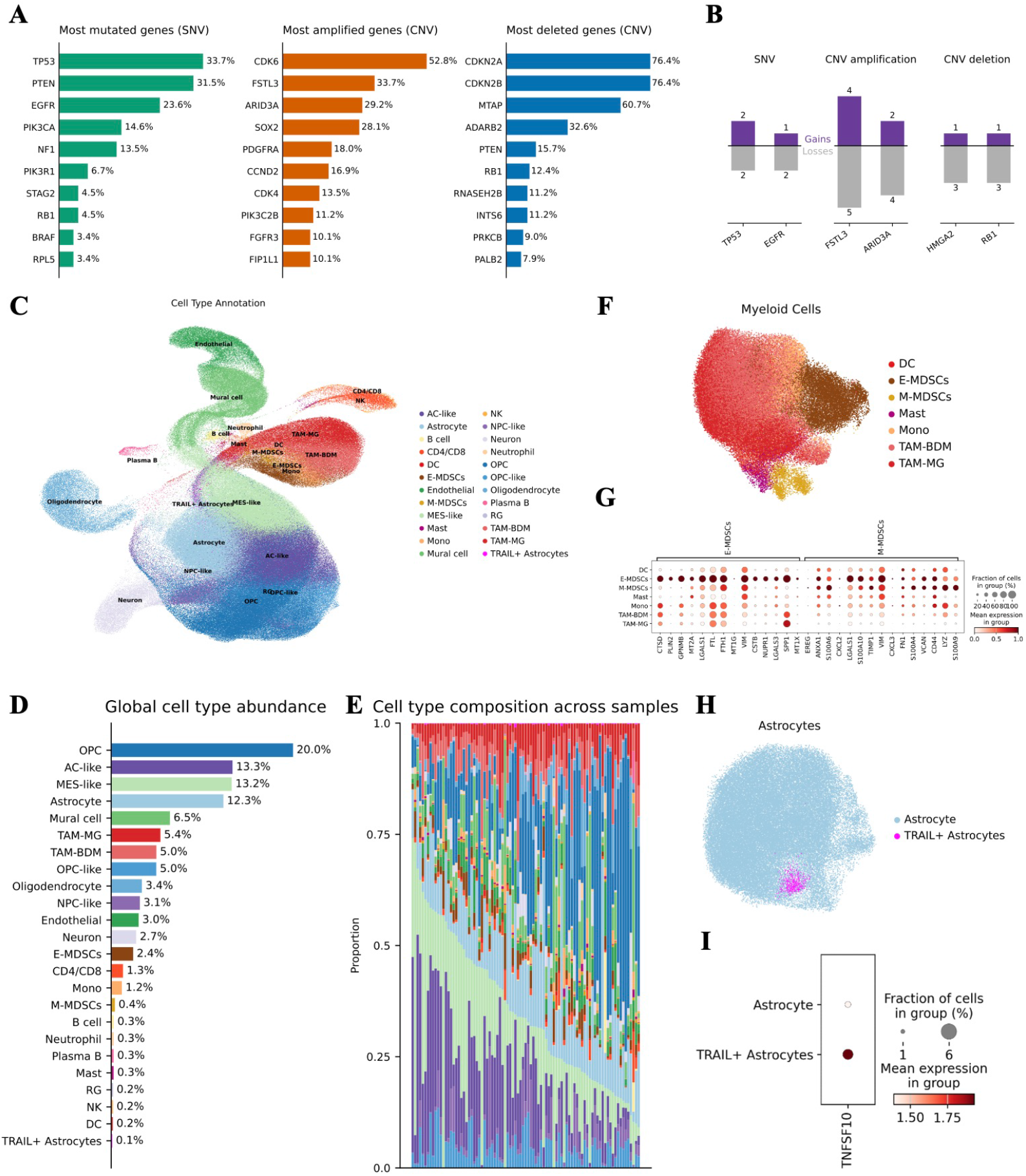
Genomic and single-cell transcriptomic landscape of the glioblastoma cohort. **A**, Summary of the genomic landscape showing the frequency of the most frequently occurring single-nucleotide variants (SNVs), copy-number amplifications, and deletions across the cohort (*n* = 89). **B**, Evolutionary dynamics of driver alterations in paired primary-relapse samples (*n* = 16), showing the gains and losses of genomic events upon tumor relapse. **C**, UMAP projection of the integrated scRNA-seq dataset annotated by major cell types, identifying malignant states (AC-like, MES-like, NPC-like, OPC-like) and diverse immune and stromal populations. **D**, Global cellular abundance across the entire cohort. **E**, Patient-specific cell type composition, highlighting significant inter-tumoral heterogeneity. **F**, Sub-clustering of the myeloid compartment reveals distinct lineages including tumor-associated macrophages (TAM; bone marrow-derived (TAM-BDM) and microglia (TAM-MG)) and myeloid-derived suppressor cells (monocytic (M-MDSCs) and granulocytic (E-MDSCs)). **G**, Dot plot of canonical marker gene expression defining the identified myeloid subpopulations. **H**, UMAP visualization of the astrocyte compartment highlighting a TRAIL+ astrocyte subpopulation. **I**, Dotplot showing expression of TNFSF10 (TRAIL) for Astrocytes as well as TRAIL+ Astrocytes.

**Figure 3:**
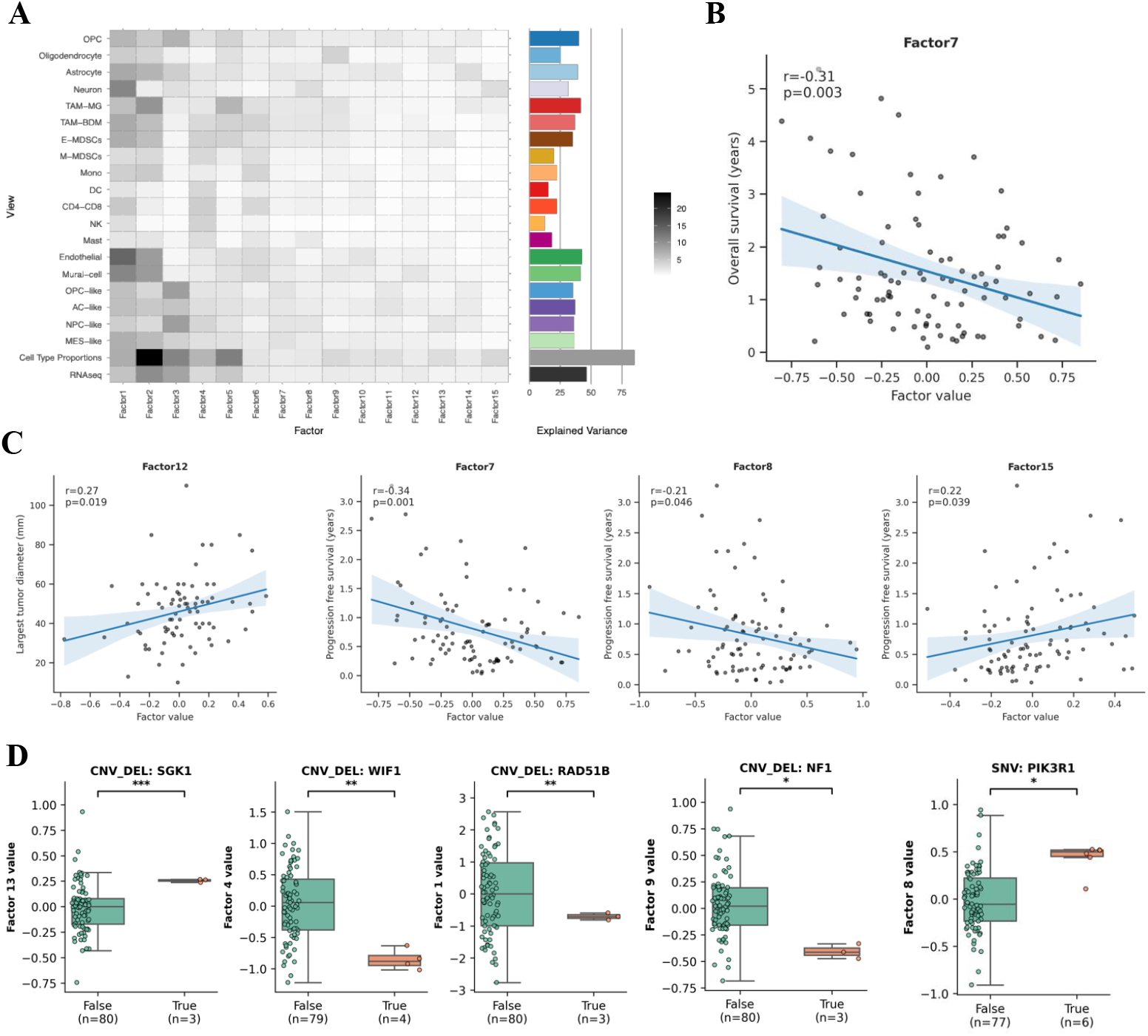
Multi-omics decomposition reveals latent factors linking genomic drivers to clinical survival and tumor morphology. **A**, Overview of MOFA model performance. The heatmap (left) illustrates the percentage of variance explained by each of the 15 latent factors (columns) across the different input views (rows). The bar plot (right) summarizes the cumulative variance explained by the model for each data view. **B**, Scatter plot showing the significant negative correlation between Factor 7 values and Overall Survival (*r* = −0.31, *p* = 0.003). **C**, Associations between latent factors and clinical covariates. Factor 12 correlates with the largest tumor diameter (*r* = 0.27, *p* = 0.019), while Factors 15 (*r* = 0.22), 7 (*r* = −0.34), and 8 (*r* = −0.21) show significant correlations with Progression-Free Survival (PFS). Shaded areas represent the 95% confidence interval of the regression estimation. **D**, Genomic characterization of latent factors. Box plots depict significant associations between specific factors and somatic alterations (Welch’s t-test, FDR-corrected). Specific factors capture copy-number deletions in *SGK1* (Factor 13), *WIF1* (Factor 4), *RAD51B* (Factor 1), and the tumor suppressor *NF1* (Factor 9), while Factor 8 is linked to SNVs in *PIK3R1*.

Next, to resolve the cellular composition of the MOSAIC glioblastoma cohort, we mapped scRNA-seq data from the tumor samples onto the GBmap reference atlas (see Methods). The resulting UMAP projection revealed a mixture of both malignant and non-malignant cell populations (Fig. 2C). Malignant cells were categorized into established transcriptomic states ^7^, including AC-like (*MLC1*), MES-like (*VIM*), NPC-like (*SOX4, DCX*), and OPC-like cells (Supplementary Fig. 1). The non-malignant compartment was dominated by a large, contiguous myeloid population alongside distinct stromal and lymphoid clusters. Global cell type abundance analysis across the cohort identified OPCs (20.0%), AC-like (13.3%), and MES-like (13.2%) cells as the most prevalent populations (Fig. 2D). However, compositional analysis across individual samples highlighted extensive inter-tumoral heterogeneity, with the relative proportions of malignant states and immune infiltrates varying significantly between patients (Fig. 2E).

Given the prominence of myeloid-mediated immunosuppression in GBM, we performed high-resolution sub-clustering of the myeloid compartment (Fig. 2F). This analysis confirmed the presence of tumor-associated macrophages of both bone marrow-derived (TAM-BDM) and microglial (TAM-MG) origin, defined by expression of *SPP1, APOE* and *CSTB* ^15^, while further resolving critical immunosuppressive populations including Monocytic and Early Myeloid-Derived Suppressor Cells (M-MDSCs and E-MDSCs). These MDSC populations, characterized by the expression of *SPP1, VCAN S1000A6*, and *S100A9* (Fig. 2G), have recently been described as exclusive features of high-grade IDH-wildtype glioblastoma ^19^.

Finally, we utilized the reference mapping to isolate the astrocyte compartment and characterize specialized functional states. This resolved a rare but distinct subpopulation of TRAIL+ astrocytes (0.1% global abundance) that clustered separately from the primary astrocyte population (Fig. 2H). In alignment with recent work identifying a glioblastoma-instructed astrocyte subset that suppresses T-cell immunity ^20^, this cluster was defined by more frequent expression of *TNFSF10* (the gene encoding TRAIL) (Fig. 2I), while otherwise sharing core astrocyte markers such as *GFAP* and *AQP4* (Supplementary Fig. 1). The specificity of *TNFSF10* to this rare cluster provides a high-confidence validation of our integrated mapping and sub-clustering approach. We also want to mention here, that like in the original publication, only ∼ 5% of the TRAIL+ Astrocytes, expressed *TNFSF10* (Supplementary Fig. 2).

The identity of all other cell populations was confirmed by the expression of cluster-specific marker genes, with the top 10 differentially expressed genes (DEGs) for each cluster detailed in Supplementary Figure 1.

Collectively, these high-resolution cellular and genomic profiles delineate a complex landscape of individual molecular layers and specialized tumor-microenvironment interactions, highlighting the extensive inter-patient heterogeneity that defines this glioblastoma cohort.

### Multi-omics integration resolves tumor heterogeneity associated with prognosis and genomic alterations

To dissect the sources of inter-patient variability, we applied MOFA to jointly model bulk RNA-seq, cell-type-specific pseudobulk expression, and cellular composition. This unsupervised integration decomposed the cohort’s heterogeneity into 15 latent factors, which captured coordinated variation across biological scales rather than being driven by single modalities (Fig. 3A). Notably, several factors accounted for a substantial fraction of the variance within the composition view, highlighting the importance of including cell type proportions as a distinct data modality.

We next assessed the clinical relevance of these latent factors by correlating factor values with patient outcomes and metadata. Factor 7 emerged as the most significant prognostic indicator, exhibiting a negative correlation with both Overall Survival (Pearson *r* = −0.31, *p* = 0.003; Fig. 3B) and Progression-Free Survival (PFS) (*r* = −0.34, *p* = 0.001; Fig. 3C). Beyond survival, the model captured morphological features, with Factor 12 showing a positive correlation with primary tumor diameter (*r* = 0.27, *p* = 0.019), effectively linking molecular profiles to macroscopic tumor growth. Additional factors, including Factor 8 (*r* = −0.21, *p* = 0.046) and Factor 15 (*r* = 0.22, *p* = 0.039), were also associated with PFS, suggesting that multi-omics integration retrieves multiple independent axes of prognostic information. Some factors also correlated with alcohol intake, smoking status as well as presence of small cells (Supplementary Fig. 5).

To determine whether these continuous factors reflect underlying somatic drivers, we tested for associations with recurrent CNVs and SNVs, restricting the analysis to the same set of GBM-related driver genes defined in the WES characterization above. This analysis revealed that specific factors act as molecular readouts for distinct genomic events (Fig. 3D). Factor 9 was significantly associated with deletions of the tumor suppressor *NF1*, capturing a known driver of the mesenchymal subtype. Similarly, Factor 8, which correlated with poor PFS, was linked to SNVs in *PIK3R1*, a key regulator of the PI3K/AKT pathway. We also identified factors associated with specific chromosomal deletions, including Factor 13 (*SGK1* deletion), Factor 4 (*WIF1* deletion), and Factor 1 (*RAD51B* deletion). Collectively, these findings demonstrate that MOFA factors not only stratify patients by clinical outcome but also reveal the downstream molecular consequences of distinct genomic drivers.

### A multicellular circuit connecting malignant progenitors and immune suppression drives poor prognosis

To elucidate the biological mechanisms driving patient outcomes, we characterized the features underlying Factor 7, the model’s strongest prognostic indicator. Feature contribution analysis identified the most prominent drivers of this factor as malignant progenitor states (NPC-like, OPC-like, MES-like, AC-like) alongside specific microenvironmental components (TAM-BDMs, Endothelial cells, OPCs, Astrocytes) (Fig. 4A). We next examined the cell–cell communication networks between these top-contributing populations to understand their interplay (Fig. 4B). This analysis revealed that malignant progenitors maintain dense signaling connections with other cell types in the microenvironment. Dissecting the functional states driving these connections, we found a fundamental divergence in pathway activity. Transcripts with high factor weights (associated with poor survival) are enriched for Coagulation and Apical Junction programs localized to NPC-like and OPC-like malignant cells (Fig. 4C). Biologically, these enrichments are consistent with aggressive tissue remodeling and invasion: Coagulation-associated genes may reflect the co-option of hemostatic factors to reorganize the extracellular matrix, while Apical Junction signaling likely points to the dynamic modulation of cell-cell adhesion required for motility. Conversely, transcripts with negative weights (associated with favorable survival) are enriched for Interferon Alpha and Gamma response within TAMs and Endothelial cells (Fig. 4C). This suggests that Factor 7 distinguishes between an immunologically active, interferon-responsive TME and a tissue-remodeling, progenitor-dominated ecosystem.

**Figure 4:**
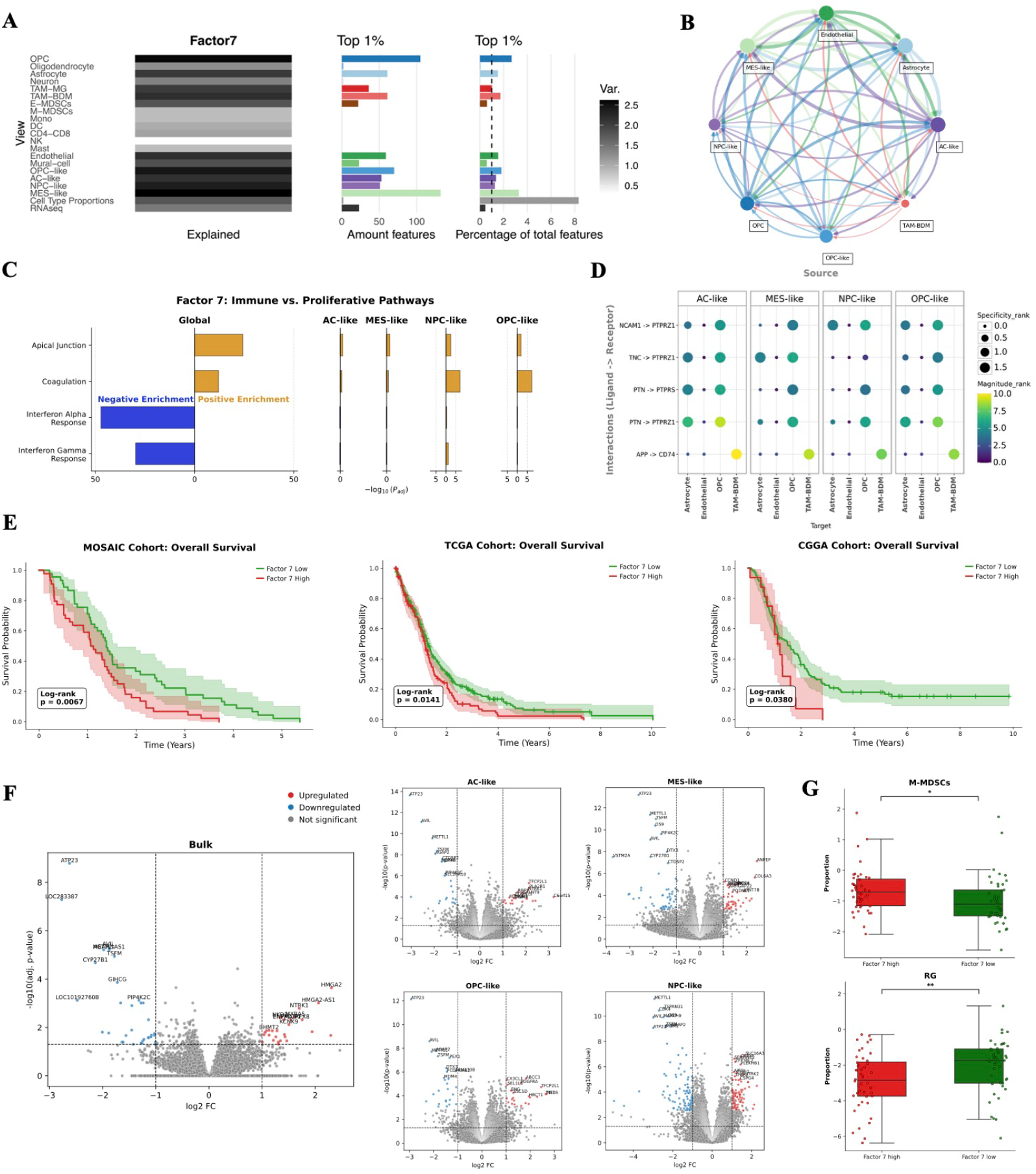
Factor 7 delineates an immunosuppressive, progenitor-driven ecosystem predictive of poor survival. **A**, The heatmap (left) displays the top weights across different views for Factor 7, while the bar plots (right) quantify the number of top features contributed by each view. **B**, Cell–cell communication network inferred from scRNA-seq data, showing interaction strength between the most contributing cell types to Factor 7. **C**, In OPC/NPC-like cells, Factor 7 positive weights (poor prognosis) enrich for Apical Junction and Coagulation; negative weights enrich for Interferon Alpha/Gamma responses. **D**, Ligand–receptor interaction analysis identifying top signaling axes. **E**, Kaplan-Meier survival analysis stratifying patients based on Factor 7 scores. The prognostic value is confirmed in the discovery cohort (MOSAIC, *p* = 0.0067) and validated in two independent cohorts using a transcriptomic proxy signature (TCGA, *p* = 0.0141; CGGA, *p* = 0.0380). **F**, Differential gene expression analysis (Volcano plots) comparing high-vs. low-Factor 7 patients in bulk RNA-seq and pseudobulked single-cell profiles. **G**, Box plots showing differential cell type abundance. The high-risk group (High Factor 7) is enriched for immunosuppressive M-MDSCs (*p <* 0.05), while the low-risk group shows higher abundance of Radial Glia (RG).

To identify the specific molecular mediators of this progenitor-driven crosstalk, we analyzed the top ligand–receptor pairs (Fig. 4D). This showed that malignant populations (AC-, MES-, NPC-, and OPC-like) are characterized by prominent signaling involving the *PTPRZ1* receptor axis, with high interaction strengths observed for the ligands *PTN, TNC*, and *NCAM1* (Fig. 4D). In addition to these tumor-intrinsic signals, we identified a APP–CD74 interaction axis linking malignant cells to myeloid TAM-BDM populations (Fig. 4D). As CD74 is a key regulator of macrophage survival and migration, this finding highlights a potential axis of communication connecting the progenitor-rich tumor compartment with the myeloid microenvironment^33,34^. Patient-level analysis confirmed coordinated expression for key ligand–receptor pairs across individual samples (see Supplementary Fig. 6).

To quantify this risk, we fitted a Cox proportional hazards model on overall survival using Factor 7 values with age, sex and MGMT-promoter methylation status as covariates. This confirmed Factor 7 as a significant risk factor for mortality (HR = 1.90, *p* = 0.04). To determine the prognostic significance of this circuit, we stratified patients in the MOSAIC cohort based on the median of Factor 7 values. This revealed a divergence in outcome, with high Factor 7 values significantly predicting shorter overall survival (log-rank *p* = 0.0067) (Fig. 4E) and progression-free survival (log-rank *p <* 0.001, Supplementary Fig. 7). Many cancer genomics data sets only comprise subsets of modalities, with RNA-seq being one of the most common data types. To enable replication of our findings in these data sets, and to facilitate the clinical translation of Factor 7 without the need for complex multi-view integration, we developed a transcriptomic proxy signature. We trained an elastic net regression model to predict Factor 7 values using only bulk RNA-seq data (Supplementary Fig. 8). The resulting sparse model selected 142 genes. Consistent with our Factor 7, this proxy signature highlighted a tissue-remodeling/migratory risk program, as evidenced by the high weighting of several key genes. Among the strongest positive predictors were *TCF21*, a bHLH transcription factor linked to mesenchymal progenitor and Epithelial-to-Mesenchymal Transition (EMT)-adjacent programs ^35^, and *FMNL3*, an actin-regulatory formin implicated in protrusion formation and invasive migration ^36^. In addition, *HYAL2*, a hyaluronidase in the hyaluronan turnover pathway, was positively weighted, consistent with established roles of hyaluronan-rich ECM remodeling in glioma invasion ^37^. Application of this signature to independent datasets confirmed the robust prognostic value of the Factor 7 proxy signature in both TCGA (*n* = 465, *p* = 0.0141) and CGGA (*n* = 133, *p* = 0.0380) cohorts (Fig. 4E), underscoring the generalizability of this risk profile across diverse ancestry patient populations.

Finally, we characterized the molecular and cellular drivers distinguishing Factor 7 high (high risk) and Factor 7 low (low risk) groups. Differential expression analysis revealed that the high-risk state is defined by a coordinated stem-like and invasive program. While bulk transcriptome analysis highlighted invasion markers including *HMGA2* and *NTRK1* (Fig. 4F), cell-type-specific profiling pinpointed the malignant compartment as the primary driver. Specifically, both NPC-like and OPC-like cells in high-risk tumors exhibited strong upregulation of the stemness factor *TFCP2L1*. This was accompanied by lineage-specific invasion programs, with *OLIG2* and *MMP17* enriched in NPC-like states, and *PDGFRA* driven by OPC-like populations. This aggressive molecular phenotype was accompanied by a compositional shift (Fig. 4G): high-risk tumors were enriched for immunosuppressive M-MDSCs (*p <* 0.05), whereas the low-risk state was associated with interferon-responsive microglia (expressing *DTX3*) and a higher abundance of Radial Glia (RG). Together, these data suggest that Factor 7 delineates a transition from a homeostatic neural precursor state to an aggressive, immune-suppressive progenitor phenotype.

### The high-risk progenitor program spatially defines the architecture of the hypoxic, perinecrotic niche

To determine the spatial organization of this molecular program, we projected Factor 7 weights onto matched Visium spatial transcriptomics sections. We calculated a spatial activity score for each spot by generating a weighted sum of the normalized expression of the factor’s top contributing genes (see Methods). In representative tumor samples, Factor 7 activity exhibited a non-random, highly structured distribution (Fig. 5A). Overlaying these molecular maps with histopathological annotations revealed that high Factor 7 activity spatially coincides with perinecrotic zones and areas of microvascular proliferation (Fig. 5B). This observation physically maps the high-risk factor to the hypoxic, semi-necrotic tissue interface.

**Figure 5:**
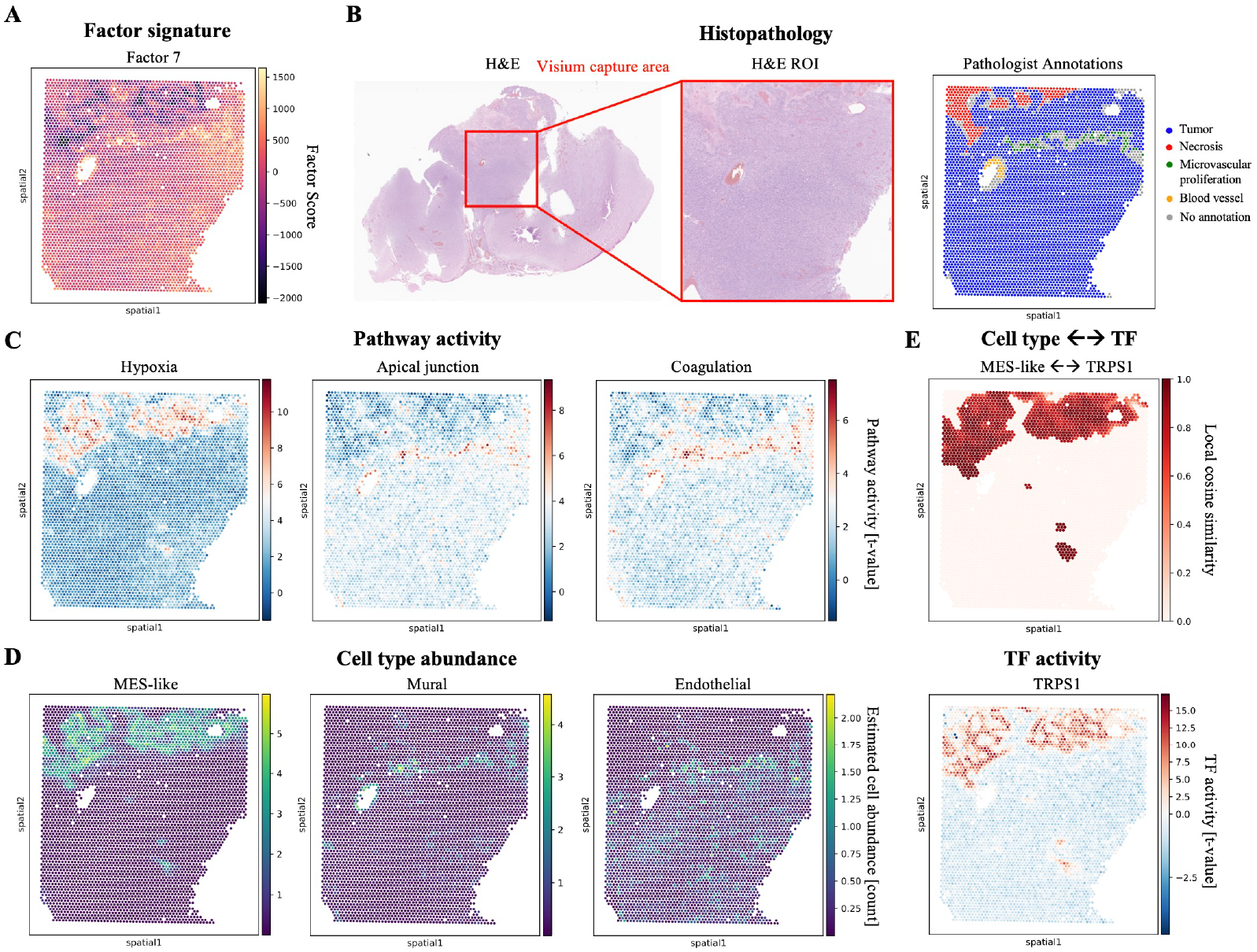
Spatial transcriptomics maps Factor 7 to hypoxic, perinecrotic niches and pinpoints regulatory drivers in situ. **A**, Spatial projection of Factor 7 activity scores. **B**, Histopathology overview displaying the Hematoxylin and Eosin (H&E) stained stained tissue with the Visium capture area (left), the H&E region of interest (middle), and spatially mapped pathologist annotations (right); Factor 7 activity scores spatially align with annotated regions of microvascular proliferation and necrosis. **C**, Spatial mapping of biological pathway activities shows strong overlap between the factor footprint, Hypoxia, and Coagulation signatures. **D**, Estimated cell type abundance maps show that MES-like tumor cells co-localize with Hypoxia in negative factor regions, while Mural and Endothelial populations align with Apical Junction and Coagulation activities in positive factor regions. **E**, Spatially resolved regulatory analysis pinpoints key transcription factors (TFs) that co-localize with malignant cells. *Top:* Local cosine similarity reveals the spatial coupling of *TRPS1* activity with MES-like tumor cells. *Bottom:* Spatial distribution of the *TRPS1* regulon.

We next dissected the biological processes spatially co-occurring with this factor. Spatial mapping revealed a strong overlap between the Factor 7 footprint and the activity patterns of Hypoxia, Coagulation, and Apical Junction signaling (Fig. 5C), consistent with the enrichments identified in the latent factor analysis. By integrating estimated cell type abundances, we observed that MES-like tumor cells co-localize with Hypoxia in negative factor regions, while Mural and Endothelial populations align with Apical Junction and Coagulation activities in positive factor regions (Fig. 5D). This segregation suggests that Factor 7 captures a polarized niche where hypoxic malignant progenitors and the vascular compartment occupy adjacent niches, providing a structural basis for the paracrine signaling circuits identified in the factor model.

Finally, we analyzed the spatial activity of transcription factors (TFs) to validate the regulatory components of this state in situ. To identify drivers specifically operating within the malignant compartment, we calculated the local cosine similarity between TF activity scores and the estimated abundance of all malignant cell states. We focused on the TF–malignant cell pair exhibiting the highest spatial co-localization. In the discovery sample, this analysis identified *TRPS1* as a key driver, showing significant spatial coupling with MES-like cells (Fig. 5E). The spatial architecture of this factor, including its localization to perinecrotic niches and the divergence of cell type neighborhoods was conserved in an independent sample, albeit driven by a distinct regulator, *NKX3-1* (Supplementary Fig. 9). This comparison highlights that while the spatial architecture of the high-risk niche is conserved, the specific transcriptional regulators driving it may exhibit inter-tumoral heterogeneity.

## Discussion

The multi-scale integration of the MOSAIC cohort presented here provides a comprehensive map of the molecular and cellular logic governing IDH-wildtype glioblastoma. By leveraging MOFA, we decomposed the extensive inter-tumoral heterogeneity of 89 patients into 15 latent factors that bridge the gap between genomic drivers and the tumor microenvironment. Our results resolve 24 distinct cell types, including rare but functionally critical populations such as TRAIL+ astrocytes and immunosuppressive MDSC subsets. The identification of these populations within our cohort provides high-resolution validation of recent models describing glioblastoma-instructed astrocytes as mediators of T-cell suppression, a phenomenon recently characterized by Faust Akl et al. ^20^, who identified an IL-11-driven astrocyte-T-cell apoptosis axis.

Factor 7 emerged as the strongest prognostic indicator in our model, delineating a transition from a homeostatic neural precursor state to an aggressive, progenitor-driven ecosystem. This axis captures a fundamental divergence in the TME: high factor weights, predictive of poor survival, are driven by Coagulation and Apical Junction programs within NPC-like and OPC-like malignant cells, whereas favorable outcomes are associated with robust Interferon Alpha and Gamma responses in the myeloid compartment. This divergence reinforces the growing consensus that the mesenchymal or progenitor phenotypes in GBM are not merely intrinsic states but are products of specific multicellular circuits ^38^. Critically, we demonstrate that the complex multi-omic architecture of Factor 7 can be distilled into a 142-gene bulk expression proxy. This signature enables the stratification of large-scale cohorts where single-cell or spatial data are unavailable, providing a cost-effective bridge to translate high-resolution discovery into routine molecular pathology. Our discovery of the *PTPRZ1* signaling axis within this progenitor-immune circuit circuit is particularly relevant, as *PTPRZ1* has recently been identified as a master regulator of glial-to-mesenchymal plasticity and a driver of invasive growth at the tumor-brain interface ^39,40^. Furthermore, our spatial analysis physically maps this identified program to the hypoxic, perinecrotic niche and areas of microvascular proliferation. This spatial coupling of malignant progenitors (MES-like cells) with Mural and Endothelial cells in semi-necrotic zones provides a structural basis for the therapy resistance observed in patients with high Factor 7 scores. This structural model is strongly supported by the recently described GRIT-Atlas ^16^, which identified a “Spatial Resistance Triad” consisting of MES-like cells, differentiation-arrested MDSCs, and collagen-secreting fibroblasts that collectively construct a fibrotic scaffold in these exact niches to shield the tumor from immunotherapy.

While the results of this study are promising, we acknowledge certain limitations. The use of pseudobulk expression for MOFA integration was a necessary trade-off to accommodate multiple modalities, which inherently masks some granular single-cell resolution. It is also important to note, that while TRAIL+ astrocytes were identified via differential expression and single-cell resolution, their low relative abundance precluded their direct inclusion as a distinct view within the MOFA model. Nevertheless, they contribute to the cell type proportion view, where Factor 11 picks them up in its top 1% feature weights. This highlights a current limitation of latent factor frameworks in capturing extremely rare cellular states that may nonetheless exert disproportionate influence on the local immune landscape. Additionally, while spatial transcriptomics provided vital in situ validation, these data were not directly incorporated as a view within the MOFA model itself. We also note that detailed histopathological annotations were available for only a subset of the cohort. Although the survival differences explained by Factor 7 were modest, the reproducibility of these findings across independent European (MOSAIC), American (TCGA), and Chinese (CGGA) cohorts confirms the robustness and potential cross-population generalizability of this marker. Given that GBM remains an essentially incurable disease with a median survival of only 12–15 months, we contend that even incremental advances in patient stratification represent a meaningful step forward.

Moving forward, the multicellular logic revealed by the MOSAIC cohort suggests several avenues for clinical translation. The development of a robust transcriptomic proxy signature offers a cost-effective tool for patient stratification in routine clinical settings where single-cell sequencing is not feasible. Future studies should focus on the longitudinal evolution of these latent factors; specifically, how the identified malignant progenitor circuit reorganizes under the selective pressure of radiotherapy and temozolomide. Given the identification of the APP-CD74, building on established CD74 roles in TAMs ^41^, and PTPRZ1 signaling axes, functional validation using patient-derived organoid (PDO) co-culture systems ^42^ will be essential to determine if disrupting these interfaces can sensitize the tumor to immunotherapy. As spatial technologies reach true subcellular resolution, directly incorporating spatial coordinates as a primary view within the MOFA framework will be critical to fully resolve the biophysical structure of the GBM niche. By defining the molecular and spatial coordinates of these aggressive niches, we provide the groundwork for targeted interventions designed to disrupt the multicellular logic that sustains this lethal cancer.

## Methods

### Patient cohort inclusion criteria

The MOSAIC dataset consists of patients with newly diagnosed, IDH-wildtype glioblastoma (median age 65.8 years; 64.0% male). Clinical profiling identified 42.3% of tumors as *MGMT* promoter methylated, with 73.9% of the cohort undergoing maximal safe resection. Integrated multi-omic profiling included matched whole-exome sequencing, bulk and single-cell RNA sequencing, spatial transcriptomics, and histopathology. For the integration via MOFA, only baseline samples and samples that had scRNAseq available were included yielding *n* = 89 samples.

For the TCGA cohort we used the `TCGA-GBM` dataset and filtered for primary tumor samples with matched RNA-seq and survival data. Additionally we removed samples that had duplicate measurements so we have independent samples.

In the CGGA cohort we utilized the `mRNAseq_693` dataset and filtered for only WHO IV grade primary tumors where survival information was available.

### Whole-Exome Sequencing and Genomic Analysis

Whole-exome sequencing was performed on the Illumina platform using a tumor-only sequencing protocol. Raw sequencing reads were processed for alignment and variant calling using the DRAGEN Bio-IT Platform (Illumina). To account for the absence of matched germline samples and to enrich for somatic alterations, identified variants were filtered against the `gnomAD v4.0` database ^43^ to exclude common germline polymorphisms. The resulting data were processed to generate binary matrices indicating the presence of somatic alterations across the cohort, focusing on two distinct classes of genomic events: single nucleotide variants (SNVs) and small insertions/deletions (indels), alongside copy number variants (CNVs).

To ensure the analysis focused on biologically relevant GBM drivers, variants were annotated and classified for potential oncogenicity using the Variant Effect Predictor (VEP) ^44^, `IntOGen` ^45^, and the `hotspots.org` database ^46^. The feature space was further restricted using established reference sets; for SNVs and indels, the analysis was limited to the set of significantly mutated genes identified in the TCGA Glioblastoma study ^27^. A gene was considered mutated if it harbored a potentially oncogenic alteration, defined as either a nonsense mutation in a known tumor suppressor gene, a frameshift indel, or a documented missense mutation at a known cancer hotspot. For copy number alterations, we restricted our analysis to a predefined set of driver genes known to exhibit significant focal events in glioblastoma. We obtained the list of significant amplifications and homozygous deletions from the TCGA Glioblastoma dataset hosted on `cBioPortal`^47,48^. Using the `GISTIC` analysis provided in this dataset, we filtered for genes with a significance threshold of *Q* ≤ 0.25. Within this set, we specifically assessed known oncogenes for amplifications and known tumor suppressor genes for homozygous deletions.

To reconstruct the evolutionary dynamics of tumor progression, we analyzed the subset of patients with matched longitudinal sequencing data (*n* = 16 pairs). Evolutionary events were categorized by comparing the binary alteration status between paired baseline (primary) and recurrence (relapse) samples. “Gains” were defined as driver alterations absent in the primary tumor but acquired at recurrence, while “Losses” were defined as alterations present in the primary tumor but undetectable in the recurrent sample.

For the Oncoprint, the gene set from the SNV gene panel was intersected with the genes set from Brennan et. al ^27^. To create the plot, the oncoPrint function of the `ComplexHeatmap` R package was used ^49^.

### Single-Cell RNA Sequencing Processing and Reference Mapping

The scRNA-seq samples were processed using `Cell Ranger v7.1.0`, followed by ambient RNA correction with `SoupX` ^50^ and doublet detection via `scDblFinder`^51^. For cell type annotation, we employed a reference mapping approach using the GBmap atlas ^15^. A `scPoli`^52^ reference model was trained on the atlas raw counts using the `annotation_level_3` labels, with `donor_id` utilized as the condition key to account for inter-individual variability. The model architecture was configured with 10 embedding dimensions and a negative binomial (NB) reconstruction loss. Training was performed for 50 epochs (including 40 pretraining epochs) with an *η* of 5, utilizing early stopping based on validation prototype loss (patience = 20) and a learning rate reduction factor of 0.1.

Query data were mapped to the reference using the `scArches` architecture surgery framework ^53^. Ambient-corrected counts were rounded to the nearest integer and aligned to the reference variable names. To ensure computational efficiency, query mapping was executed in batches of 10 donors. Each batch underwent fine-tuning for 50 epochs (40 pretraining) with an *η* of 10. Final cell type assignments were generated through the `scPoli` classification module, which transferred the `annotation_level_3` labels from the reference latent space to the query cells.

For the downstream analysis of the myeloid compartment, `scanpy`^54^ was utilized. Cells with a reference mapping uncertainty higher than 0.7 were excluded. We specifically isolated the TAM-MG, TAM-BDM, Mono, DC, and Mast cell populations, as identified by the reference mapping, restricting this sub-analysis to baseline samples. To identify M-MDSCs and E-MDSCs, we first identified the top 2,000 highly variable genes, accounting for batch effects by using `sample_id` as the batch key. Principal Component Analysis (PCA) was performed on these genes, and the resulting embeddings were integrated using the `Harmony` algorithm ^55^ to mitigate sample-specific technical variation.

Following integration, a neighborhood graph was constructed based on the Harmony-corrected principal components. We then performed manifold embedding using Uniform Manifold Approximation and Projection (UMAP) ^56^ and identified distinct cell clusters via Leiden clustering ^57^ at a resolution of 0.5. This high-resolution clustering of the myeloid lineage allowed for the specific annotation and segregation of MDSC subsets. To identify TRAIL+ astrocytes, we isolated cells annotated as “Astrocyte” and applied the same computational pipeline and parameters used for the myeloid analysis. Following Harmony-based integration and Leiden clustering (*resolution* = 0.5), we specifically identified a TRAIL+ astrocytes based on marker expression *TNFSF10*, while the remaining cells were maintained as a single astrocyte population for downstream comparisons.

### Multi-Omics Factor Analysis (MOFA) Integration

For the bulk RNA-seq analysis, raw Ensembl IDs were mapped to gene symbols and biotypes using the `MyGene.info` database ^58^. To ensure a high-quality protein-coding focused dataset, we filtered the transcriptome by excluding non-coding RNA species (snRNA, snoRNA, rRNA, scRNA, and tRNA), pseudogenes, and genes with “pseudogene” in their metadata. Expression counts for duplicate symbols were aggregated by summation, and any remaining unmapped Ensembl IDs were removed.

Data quality control was performed by filtering out genes expressed in fewer than 50% of the samples. To identify biologically informative features, we performed variance-stabilizing transformation using a log_2_(count + 1) scale and implemented a Seurat-inspired highly variable gene (HVG) selection workflow. Specifically, genes were partitioned into 20 bins based on their mean expression levels. Within each bin, a Z-score for variance was calculated to identify genes with high dispersion relative to their expression magnitude. The top 5000 genes with the highest variance Z-scores were selected for downstream analysis. Finally, sample identifiers were standardized by removing underscores and ‘mRNA’ suffixes to ensure consistency across the study datasets.

Cell type proportions were generated by grouping the annotated single-cell dataset by `sample_id` and `cluster_id`. To account for the compositional nature of single-cell proportion data and the presence of zero counts, we utilized the `zCompositions`^59^ R package. Specifically, zero values were imputed using the Bayesian-multiplicative replacement method via the `cmultRepl` function to produce a complete proportion matrix. The resulting proportions were then subjected to a Centered Log-Ratio (CLR) transformation to move the data from a constrained simplex to an unconstrained Euclidean space, mitigating the dependency between cell type fractions. The transformation was calculated for each sample as the logarithm of the ratio between each cell type proportion and the geometric mean of all cell type proportions in that sample. These CLR-transformed values were utilized as the final feature set for downstream multi-omic integration.

Multi-omic integration was performed using MOFA+^22^, following the code and tutorial described by Pekayvaz et al. ^24^. The analysis was restricted to baseline samples, excluding two individuals for whom scRNA-seq data were unavailable, resulting in a final cohort of *n* = 89 samples. To ensure robust factor inference, low-abundance cell types, specifically RG, Plasma B cells, B cells, Neutrophils, and TRAIL+ Astrocytes, were excluded from the pseudobulking process due to sparsity across samples.

For the scRNA-seq views, feature selection was performed using two filtering criteria: genes were retained if they were expressed in at least 1% of cells in at least one sample, or in 5% of cells across at least two samples. Both scRNA-seq and bulk RNA-seq views underwent library size adjustment and log transformation, followed by feature-wise and sample-wise quantile normalization. For the single-cell views, the feature set was further refined to the top 30% highly variable genes. During model training, views were scaled and weighted according to their respective feature dimensions, and the final MOFA model was configured to extract 15 latent factors.

Functional characterization of the MOFA factors was performed using the `run_enrichment_pathway` function. This enrichment analysis implements a methodology adapted from the Principal Component Gene Set Enrichment (PCGSE) framework ^60^. We utilized the Cancer Hallmark gene sets from the Molecular Signatures Database (`MSigDB`) ^61^ as the reference pathway collection. To ensure the reliability of the enrichment scores, a coverage threshold of 0.2 was applied, requiring that at least 20% of the genes within a given pathway were present among the features of the respective factors. Significant pathways were identified based on their association with the latent factors, providing biological context for the inferred axes of variation.

To characterize the clinical relevance of the inferred latent factors, we performed a systematic correlation analysis between factor values and available patient metadata. For continuous variables, including overall survival years, progression-free survival years, and tumor diameter, we calculated Pearson correlation coefficients (*r*) and associated p-values, visualized through linear regression models. For categorical clinical annotations, such as necrosis, smoking status, and specific histopathological features, we assessed differences in factor distributions across groups using independent two-sample t-tests (for binary comparisons) or one-way Analysis of Variance (ANOVA, for multiclass comparisons). Statistical significance was categorized and annotated on box-and-whisker plots (∗ < 0.05, ∗∗ < 0.01, ∗∗∗ < 0.001). This approach allowed us to identify specific axes of multi-omic variation that were significantly associated with disease progression and patient lifestyle factors.

Associations between MOFA factors and genomic features, including SNVs and CNVs, were evaluated by comparing factor distributions across mutated and wild-type groups. For each gene-factor pair, a filtering threshold was applied requiring at least *n* = 3 samples in both the mutated and wild-type groups before statistical testing. We employed independent Welch’s t-tests to assess statistical differences, with p-values adjusted for multiple testing using the Benjamini-Hochberg False Discovery Rate (FDR) procedure. Significant associations were visualized through box-and-whisker plots.

### Cell-Cell Communication Inference

To systematically map the intercellular signaling networks driving tumor progression, we performed ligand-receptor (L-R) interaction analysis using the `LIANA` (Ligand-Receptor Analysis) framework ^62,63^. Single-cell RNA sequencing data were first normalized (scaling target sum to 10^4^) and log-transformed (log(*x* + 1)). To ensure robust interaction calling, we employed a consensus approach by aggregating predictions from six diverse methods: `CellPhoneDB` ^64^, `Connectome` ^65^, `LogFC`, `NATMI` ^66^, `SingleCellSignalR`^67^, and `Geometric Mean`.

Interactions were aggregated by cell type, and a consensus rank was calculated based on the magnitude and specificity of expression. To focus on paracrine signaling relevant to the tumor microenvironment, we filtered out autocrine interactions (where source and target cell types are identical). Subsequent analysis focused specifically on the top-contributing cell types identified by Factor 7 (malignant progenitors, endothelial cells, and myeloid populations). Interactions were prioritized based on interaction magnitude (reflecting signal strength) and specificity (reflecting the exclusivity of the signal to a given pair).

To assess the stability of these intercellular networks across the patient cohort, we performed a sample-level correlation analysis on top-ranked candidate pairs. Interactions were selected to cover the primary signaling axes linking the malignant compartment (AC-like, NPC-like, MES-like, and OPC-like states) with the tumor microenvironment (myeloid and endothelial cells). For each selected ligand-receptor pair, we aggregated single-cell expression data to the sample level by calculating the mean expression of the ligand in the source cell population and the receptor in the target cell population for each patient (*n* = 89). We then computed the Pearson correlation coefficient (*R*) between these matched sample-level aggregates to quantify the extent to which ligand abundance in the sender population predicted receptor expression in the receiver population across the cohort.

### Patient Stratification and construction of the transcriptomic proxy signature

Patients in the discovery cohort were stratified based on the median Factor 7 value (−0.0068), defining Factor 7 High and Factor 7 Low groups. To identify molecular signatures associated with this stratification, we performed differential expression (DE) analysis on the bulk RNA-seq data using `PyDESeq2`^68^. The model utilized these groups as design factors, with the Factor 7 Low group serving as the reference level. We incorporated Cook’s distance refitting to handle outliers and adjusted p-values using the Benjamini-Hochberg procedure. Significance was defined by an adjusted p-value (*P*_*adj*_) < 0.05 and an absolute log2 fold change (|log2FC|) *>* 1.

In parallel, cell-type-specific differential expression was conducted using a pseudo-bulk approach. Single-cell counts were aggregated by sample id and `cluster_id` using the `Decoupler`^69^ package. To ensure statistical robustness, we excluded pseudo-bulk samples with fewer than 10 cells or 100 total counts, and only cell types represented in at least three samples per group were retained. DE testing was then performed for each eligible cell type using the `PyDESeq2` framework, consistently using the Factor 7 Low group as the reference. Resulting differential features were visualized via volcano plots, with top-ranking genes identified by their statistical significance and magnitude of fold change.

To assess differences in cellular composition between the Factor 7 High and Factor 7 Low groups, we compared the CLR-transformed cell type proportions using independent two-sample t-tests. Statistical significance was determined at an alpha level of 0.05, with resulting p-values visualized as significance stars (∗ < 0.05, ∗ ∗< 0.01, ∗ ∗ ∗ < 0.001) above grouped box-and-whisker plots. These visualizations were generated to highlight the relative expansion or contraction of specific cell lineages, with individual sample distributions overlaid as jittered points to ensure transparency of the underlying data variance.

To enable the translation of our multi-omic findings to independent cohorts where only transcriptomic data is available, we developed a gene-expression-based proxy for Factor 7. Raw counts from the discovery cohort and the TCGA and CGGA validation cohorts were normalized to log_2_(CPM + 1) to ensure cross-platform comparability. We identified a set of overlapping genes across all datasets to serve as the feature space.

A predictive model was constructed using ElasticNet regression within the `scikit-learn`^70^ framework, targeting the Factor 7 latent values as the continuous dependent variable. The discovery dataset was split into training (80%) and internal testing (20%) sets. Features were standardized using a `StandardScaler` fit exclusively on the training data to prevent data leakage. We employed 5-fold cross-validation (`ElasticNetCV`) to optimize the regularization parameters, testing a range of L1 ratios [0.1, 0.5, 0.7, 0.9, 0.95, 0.99, 1.0]. The final model, utilizing the optimal alpha (0.04148) and L1 ratio (0.10), identified a parsimonious gene signature for Factor 7 prediction.

For external validation in the TCGA and CGGA cohorts, predicted Factor 7 values were calculated and patients were stratified into high- and low-risk groups based on the discovery cohort’s median threshold.

### Survival Modeling

Survival analysis was performed using Overall Survival and Progression-Free Survival as primary endpoints. To evaluate the independent prognostic value of Factor 7, we employed multivariate Cox Proportional Hazards (CoxPH) models using the `lifelines`^71^ Python library. The models were adjusted for relevant clinical covariates, including age at diagnosis, sex, and *MGMT* promoter methylation status. Categorical variables (sex and *MGMT* status) were transformed into indicator variables using one-hot encoding prior to model fitting.

Survival distributions for the Factor 7 stratified groups were estimated using Kaplan-Meier (KM) curves. The statistical significance of differences between survival curves was assessed using the log-rank test for both the discovery (multivariate log-rank) and validation (TCGA and CGGA) cohorts. For the validation cohorts, survival analysis was based on the predicted Factor 7 clusters generated by the ElasticNet proxy signature. All KM plots include 95% confidence intervals, and p-values < 0.05 were considered statistically significant.

### Spatial Transcriptomics and Histopathology Integration

To spatially resolve the molecular programs identified by the multi-omics integration, we utilized 10x Genomics Visium spatial transcriptomics data with matched H&E histology. High-resolution H&E images were aligned to the Visium capture area using tissue positions, allowing for the direct overlay of molecular data with morphological features. Raw sequencing data were processed using the 10x Genomics `Space Ranger` pipeline ^72,73^, which performed image alignment, tissue detection, and gene counting. The resulting datasets consisted of arrays containing up to 5,000 spots across 16,927 genes; for all downstream analyses, we utilized the filtered subset of spots identified as being “under-tissue” by the `Space Ranger` software. For a subset of samples, regions of interest, including necrosis, microvascular proliferation, and tumor core, were manually annotated by a neuropathologist to validate the spatial localization of computational signatures.

To project the latent multi-omic factors onto the spatial tissue, we calculated spatial activity scores. Visium gene expression counts were first library-size normalized, log-transformed, and quantile-normalized across spots to ensure robust comparability. We then computed a weighted sum of the normalized spatial expression data using the gene weights extracted from the relevant MOFA factor (e.g., Factor 7). This approach allowed us to visualize the spatial distribution of the high-risk progenitor program in situ.

Biological pathway activities were inferred spatially using `decoupler`^69^. We utilized the `PROGENy` database ^74^ to estimate the activity of canonical signaling pathways (e.g., Hypoxia, Coagulation, JAK-STAT) at single-spot resolution via a multivariate linear model. To map the cellular architecture, we performed spatial deconvolution using `Cell2Location`^75^. A reference regression model was first trained on the cohort’s single-cell RNA-seq data to characterize the expression signatures of the 24 identified cell types. These signatures were then used to estimate the abundance of each cell type at every spatial location (*N*_*cellsperlocation*_ = 5).

Finally, to identify the transcriptional regulators driving these spatial niches, we performed a spatially resolved regulatory analysis. Transcription factor (TF) activities were estimated using the `CollecTRI` gene regulatory network ^76^ via a univariate linear model. To link these regulators to specific cell populations, we computed the local spatial correlation between TF activity and estimated cell type abundance using the `bivariate` function in the `LIANA` framework ^63^. Local associations were quantified using a Gaussian kernel (bandwidth=200), and TF-cell type pairs were ranked based on their spatially weighted cosine similarity. This metric was used to prioritize candidate regulators that exhibited strong spatial coupling with malignant cell states for downstream characterization.

## Data availability

TCGA data can be downloaded at https://www.cancer.gov/tcga. The CGGA data is available at https://www.cgga.org.cn/download.jsp. MOSAIC data is not publicly available; however, access to a representative subset of 60 patients may be requested via MOSAIC-Window (https://www.mosaic-research.com/mosaic-window).

## Code availability

The code will be made public upon publication of the manuscript.

## Acknowledgements

K.T. is supported by the Helmholtz Association under the joint research school “Munich School for Data Science—MUDS”. J.T. acknowledges support from the Gates Cambridge Trust via the Gates Cambridge Scholarship. M.H. is supported by the Chan Zuckerberg Foundation (2019-202666, 2021-237882) and the DZHK (German Center for Cardiovascular Research) projects 81Z0600106 and 81Z0600105. G.S.K.S acknowledges funding from the Wellcome Trust (065807/Z/01/Z) (203249/Z/16/Z), the UK Medical Research Council (MRC) (MR/K02292X/1), ARUK (ARUK-PG013-14), Michael J Fox Foundation (16238; 022159), and Infinitus China Ltd. This study makes use of data generated by the MOSAIC consortium (Owkin, Charité – Universitätsmedizin Berlin, Lausanne University Hospital - CHUV, Erlangen Hospital, Gustave Roussy Institute, University of Pittsburgh) and made available through the GBM Hackathon organized by Owkin, as satellite event of the 2025’ Paris AI Action Summit. Readers should note that the MOSAIC consortium bears no responsibility for the further analysis or interpretation of these data beyond what published by the MOSAIC consortium partners. The results shown here are in part based upon data generated by the TCGA Research Network: https://www.cancer.gov/tcga.

