## Supplementary Figures for "Spatially Integrated Multi-Omics reveals the Multicellular Landscape of Progenitor-Driven Glioblastoma Progression"

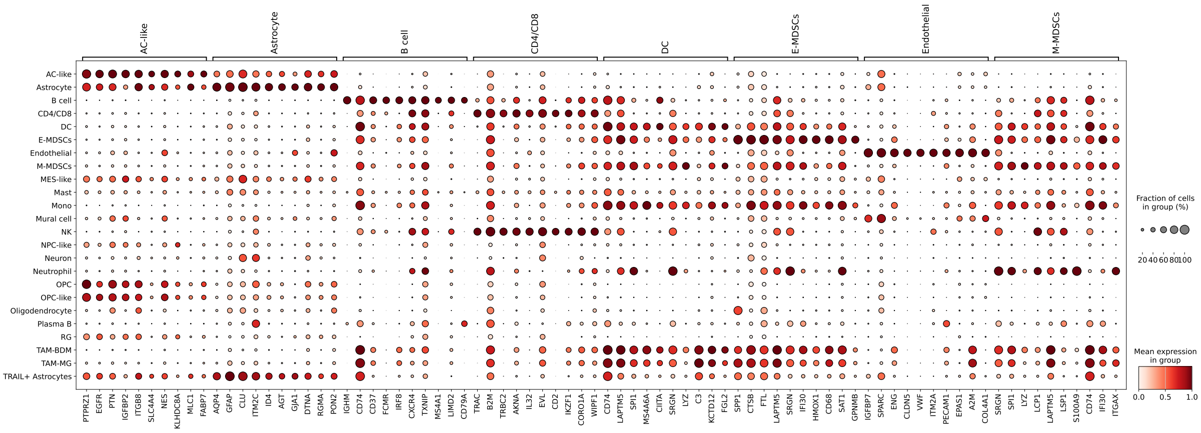

(a) Expression markers for first gene group

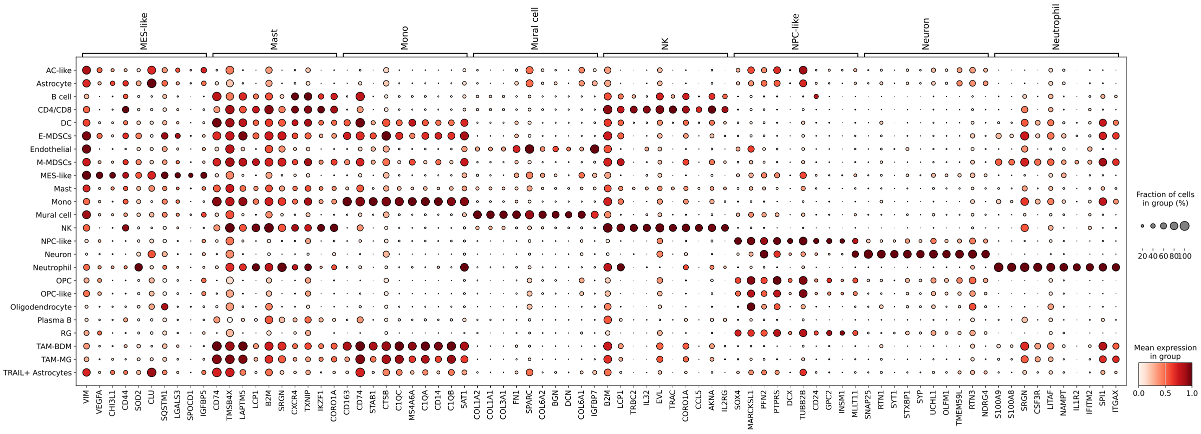

(b) Expression markers for second gene group

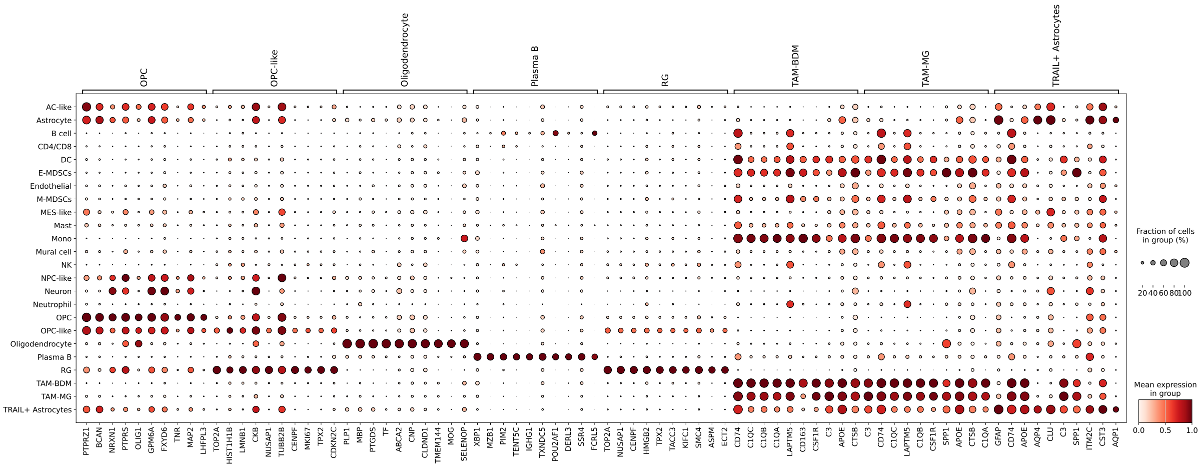

(c) Expression markers for third gene group

Supplementary Figure 1: **Top 10 differentially expressed genes identifying specific cell clusters.** The dot plots illustrate the top 10 marker genes identified for each cell type cluster.

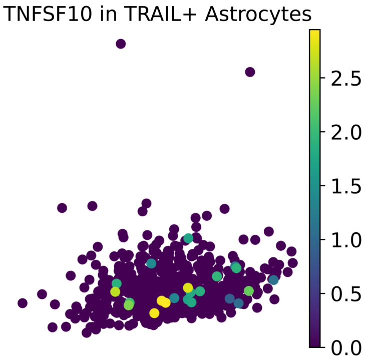


Supplementary Figure 2: **Restricted expression of *TNFSF10* in the TRAIL+ astrocyte subpopulation.** The feature plot displays the specific expression of *TNFSF10* (encoding TRAIL) within the astrocyte compartment. While this marker defines the distinct TRAIL+ cluster, expression is sparse and detected in approximately 5% of the cells within this specific subpopulation, consistent with previously reported frequencies.

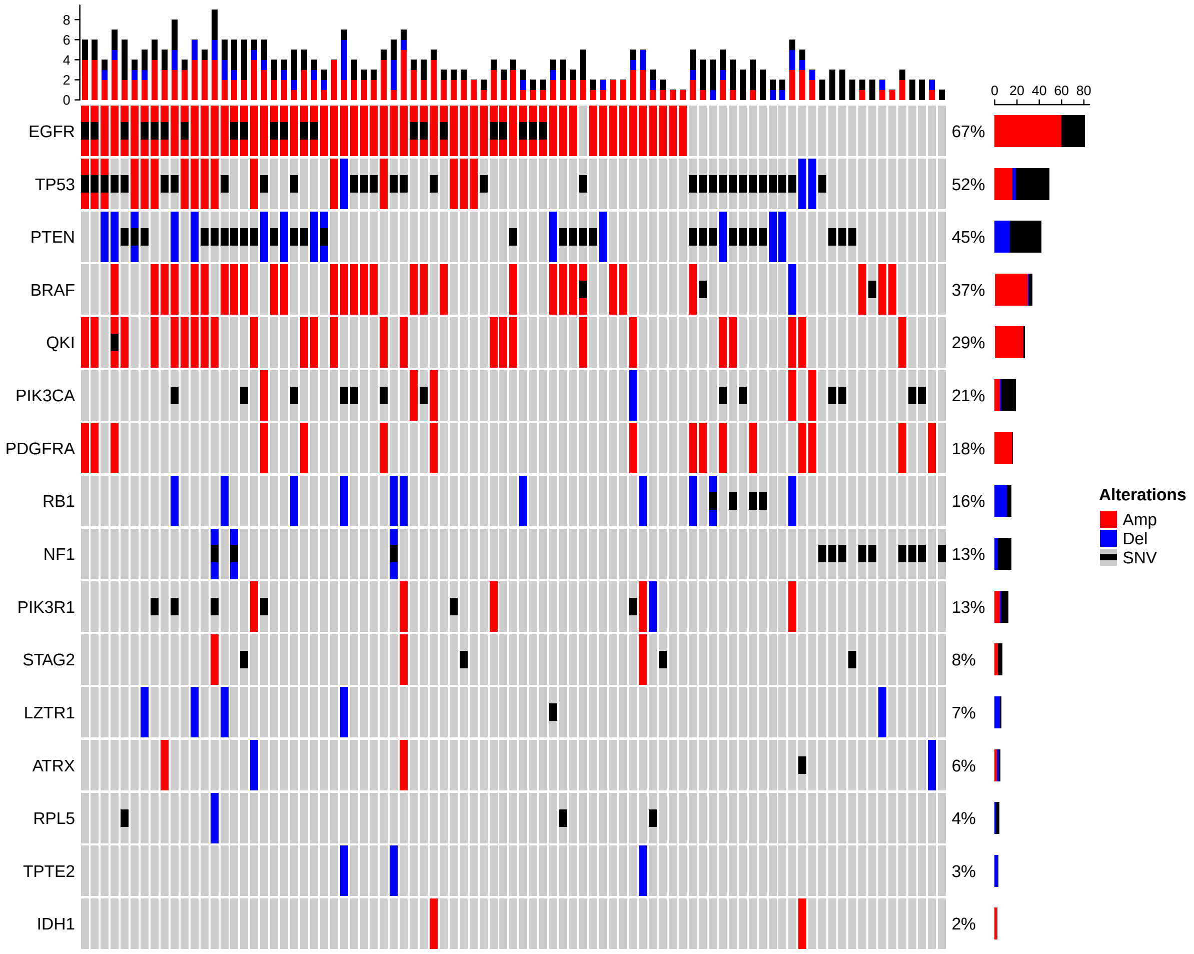


Supplementary Figure 3: **Landscape of somatic alterations in the MOSAIC glioblastoma cohort.** The oncoprint summarizes the genetic alterations (SNVs and CNVs) across the cohort ($n = 89$), restricted to the set of significantly mutated genes identified in the TCGA Glioblastoma study. This visualization highlights *EGFR* as the most frequently altered gene (67% of samples) when combining mutations and copy-number changes.

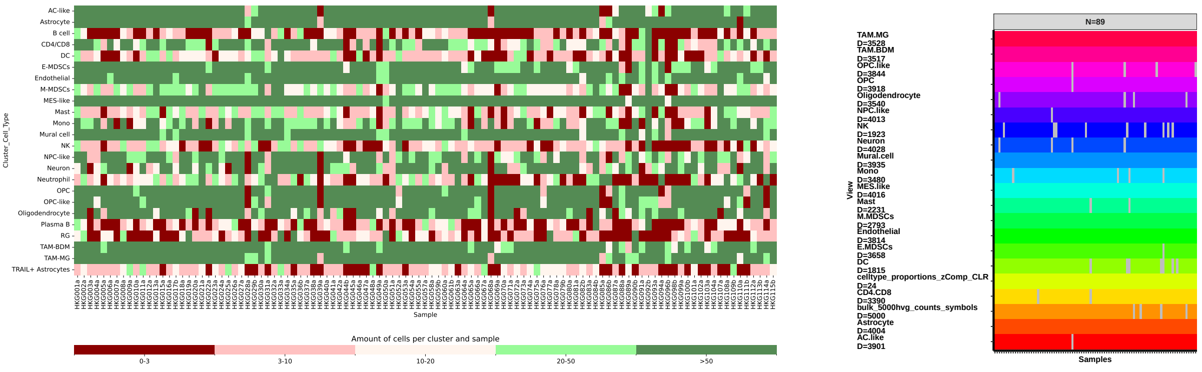


(a) (b)

Supplementary Figure 4: **Cellular composition heterogeneity and multi-omics integration strategy.** (a) Heatmap displaying the absolute abundance of cells per cell type (rows) across individual patient samples (columns) in the MOSAIC cohort. Color intensity represents binned cell counts, highlighting the inter-tumoral heterogeneity in tissue composition. (b) Overview of the input views integrated into the MOFA model after filtering. The model combines bulk RNA-seq (top 5000 highly variable genes), cell type proportions, and cell-type-specific pseudobulk expression profiles for sufficiently abundant populations. Blank boxes indicate that this view is not available for the sample.

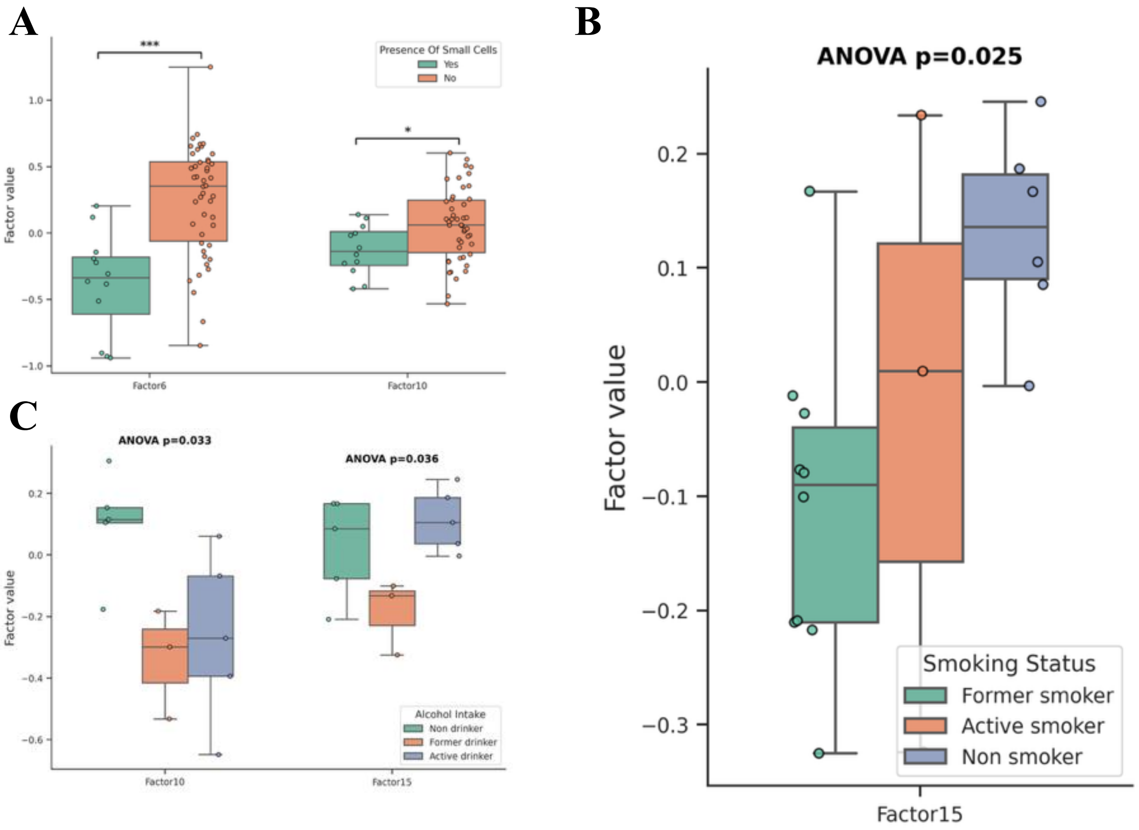


Supplementary Figure 5: **Associations of latent factors with categorical clinical data.** Box plots depict significant associations between specific factors and **A** Presence of small cells (Welch's t-test,), **B** smoking status (ANOVA test), and **C** alcohol intake (ANOVA test).

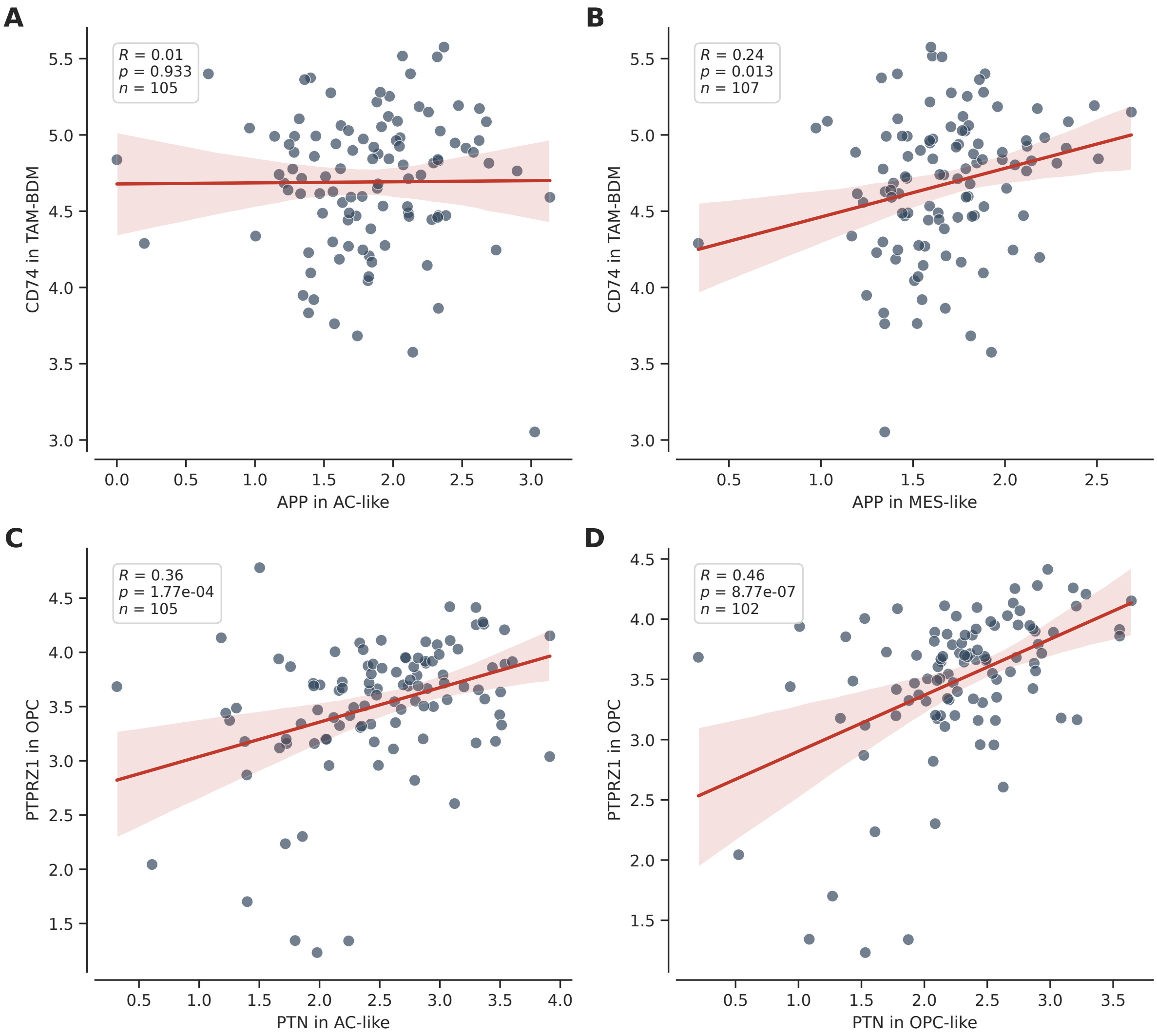


Supplementary Figure 6: **Correlation of ligand–receptor expression levels across patient samples.** Scatter plots displaying the relationship between the mean log-normalized expression of ligands in source cell types (x-axis) and receptors in target cell types (y-axis). Each dot represents an individual patient sample ($n$ indicates sample size). The red line represents the linear regression fit with the 95% confidence interval (shaded region). Pearson correlation coefficients ($R$) and $p$-values are annotated for each interaction pair. **A-B**, Analysis of the *APP–CD74* axis between malignant progenitors and TAM-BDMs, showing *APP* expression in AC-like cells (**A**) and MES-like cells (**B**). **C-D**, Analysis of the *PTN–PTPRZ1* axis between malignant progenitors and OPCs, showing *PTN* expression in AC-like cells (**C**) and OPC-like cells (**D**).

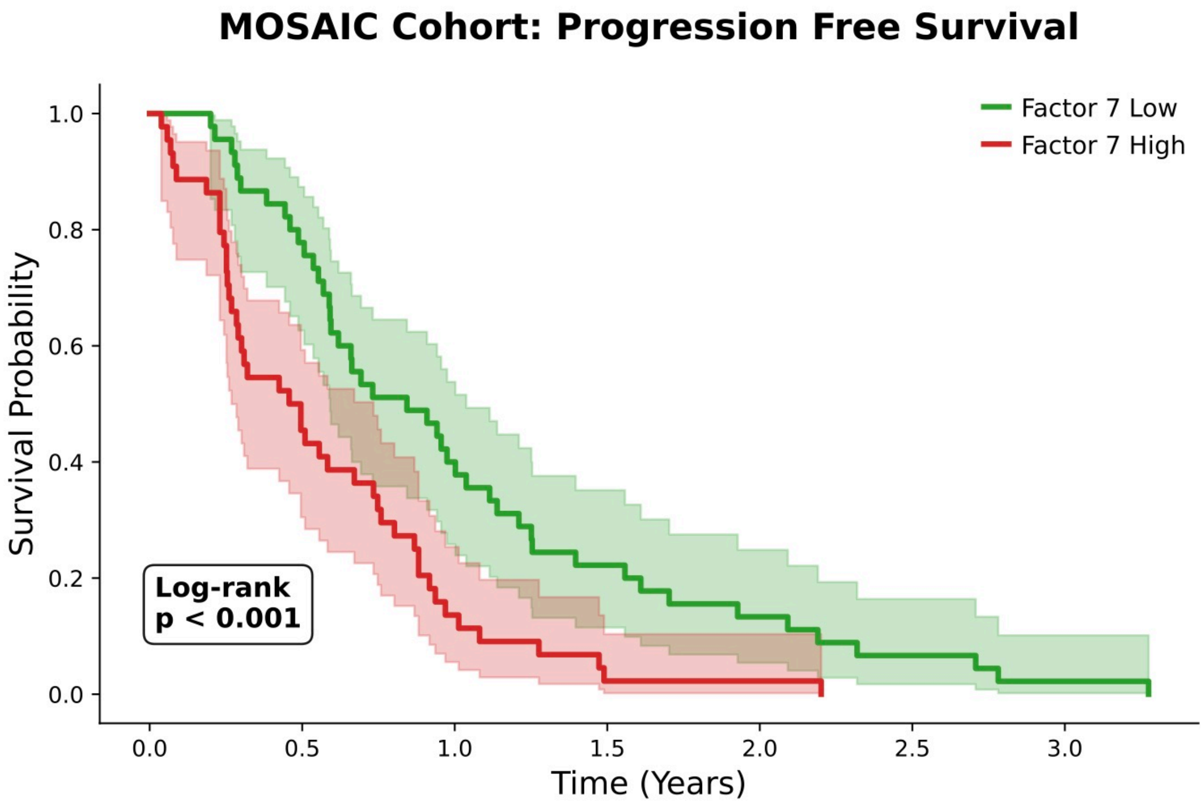


Supplementary Figure 7: **Prognostic value of Factor 7 for progression-free survival.** The Kaplan-Meier survival curve stratifies patients from the MOSAIC cohort into high and low risk groups based on the median Factor 7 score. Patients with high Factor 7 scores exhibit significantly shorter progression-free survival (PFS) compared to the low scoring group (log-rank test $p < 0.001$), confirming the factor's association with aggressive disease trajectories.

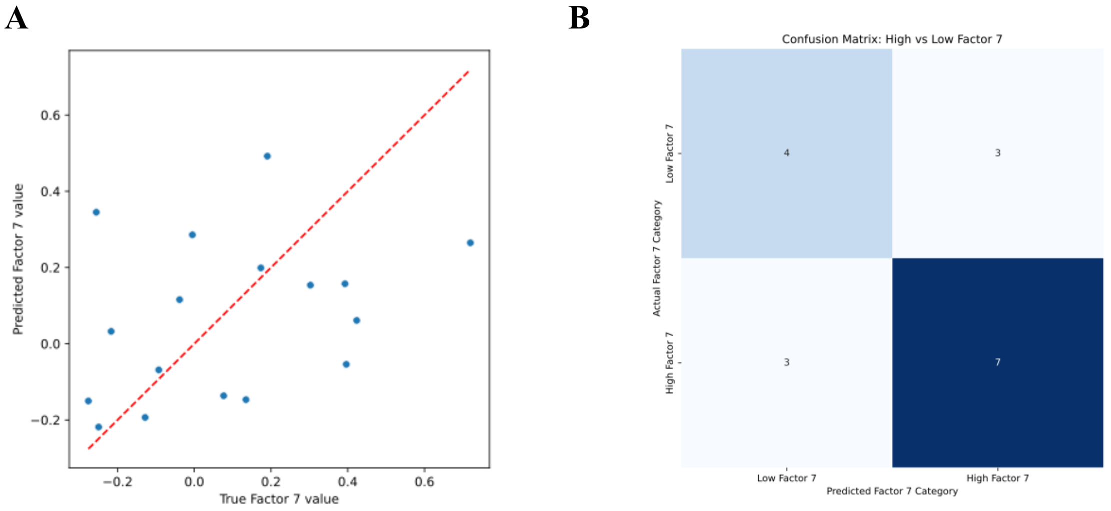


Supplementary Figure 8: **Validation of the elastic net proxy signature for Factor 7. A** depicts the true and predicted factor 7 values, while **B** shows the confusion matrix when split into High and Low Factor 7 groups based on the median in the discovery cohort.

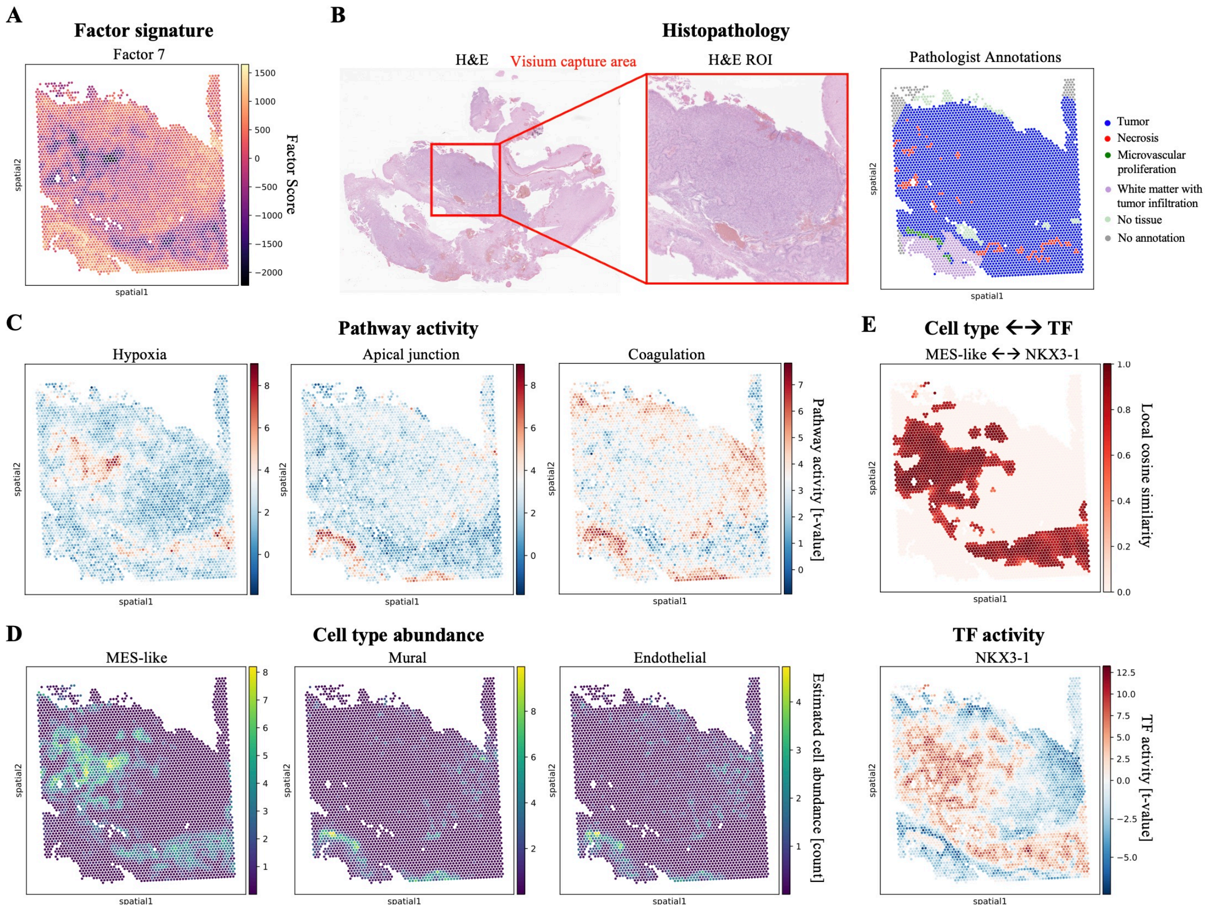


Supplementary Figure 9: **Validation of Factor 7 spatial architecture in an independent sample.** **A**, Spatial projection of Factor 7 activity scores. **B**, Histopathology overview displaying the H&E stained tissue with the Visium capture area (left), the H&E region of interest (middle), and spatially mapped pathologist annotations (right); Factor 7 activity scores spatially align with annotated regions of microvascular proliferation and necrosis. **C**, Spatial mapping of biological pathway activities shows strong overlap between the factor footprint, Hypoxia, and Coagulation signatures. **D**, Estimated cell type abundance maps show that MES-like tumor cells co-localize with Hypoxia in negative factor regions, while Mural and Endothelial populations align with Apical Junction and Coagulation activities in positive factor regions. **E**, Spatially resolved regulatory analysis pinpoints key transcription factors that co-localize with malignant cells. *Top:* Local cosine similarity reveals the spatial coupling of *NKX3-1* activity with MES-like tumor cells. *Bottom:* Spatial distribution of the *NKX3-1* regulon.
